# Loss of function of LprG-Rv1410c and MmpL11 homologues in *Mycobacterium smegmatis* leads to altered glycopeptidolipid profile and decreased cellular c-di-GMP levels during biofilm formation

**DOI:** 10.64898/2026.09.10.750705

**Authors:** Mariah Desroches, Miranda Coldren, Lisa-Marie Nisbett

## Abstract

The cell envelope of many bacteria possesses components that play central roles in bacterial pathogenesis. Unique to mycobacteria is the presence of a lipid-rich cell wall which is critical for the virulence of pathogenic mycobacteria such as *Mycobacterium tuberculosis* (*Mtb*). The biosynthesis of mycobacterial cell wall lipids has been well characterized, but the mechanisms of lipid transport still remain largely unknown. MmpL (mycobacterial membrane protein large) proteins have been implicated in the biosynthesis and/or transport of mycobacterial cell wall lipids, but due to their cellular location, it is still unclear how cell wall lipids are transported beyond the inner membrane. Here, we further investigate the role of two conserved lipid transport pathways LprG-Rv1410c and MmpL11 in cell envelope biogenesis during biofilm formation in *Mycobacterium smegmatis*. We found that deletion of the *lprG*-*rv1410c* operon homologues *MSMEG_3070-3069* and *mmpL11* (*MSMEG_0241*) simultaneously led to similar biofilm defects as observed in the *MSMEG_3070-3069* and *mmpL11* mutants. Analysis of pellicle biofilms, total lipid extracts, gene expression and cellular c-di-GMP levels revealed that the shared biofilm defect is directly correlated with significant decreases in cellular c-di-GMP levels, but may also be due to changes in the synthesis and/or localization of glycopeptidolipids (GPLs), and changes in the expression of GPL biosynthesis and transport genes. Our findings therefore suggest that while both LprG-Rv1410c and MmpL11 pathways are involved in modulating cellular c-di-GMP levels during biofilm formation, only LprG-Rv1410c are important for regulating GPL synthesis and/or surface localization, and MmpL11 may play a broad role in fine-tuning GPL levels.

**IMPORTANCE:** The lipid-rich cell wall has been demonstrated to be essential for the virulence of pathogenic mycobacteria such as *Mycobacterium tuberculosis* (*Mtb*). The biosynthesis of mycobacterial cell wall lipids has been well characterized, but the mechanisms of lipid transport still remain largely unknown. Here, we demonstrate that LprG-Rv1410c and MmpL11 impact biofilm formation in *M. smegmatis* via modulation of GPL biosynthesis and/or cell wall localization and cellular c-di-GMP levels. This work further underscores the importance of lipid transport pathways in cell wall biogenesis and provides additional insights into how GPL synthesis and/or cell wall transport may be regulated during mycobacterial biofilm formation, with implications for better understanding the connection between mycobacterial cell wall biogenesis, biofilm formation and mycobacterial physiology overall.

## INTRODUCTION

Tuberculosis (TB) still remains a major global health threat; in 2024 approximately ten million people fell ill with TB and approximately one million died from the disease (1). Further, the continued emergence and spread of multi-drug resistant TB highlights the urgent need for novel therapeutic approaches against *Mycobacterium tuberculosis* (*Mtb*), the causative agent of TB in humans. The drug resistance of *Mtb* is due in part to a well-armored cell wall which is complex and unique in composition. The outer membrane (commonly referred to as the mycomembrane) is composed of an inner leaflet rich in mycolic acids that are covalently attached to an underlying arabinogalactan-peptidoglycan layer, and an outer leaflet composed of diverse noncovalently associated free lipids such as trehalose 6,6’-dimycolate (TDM), phthiocerol dimycocerosate (PDIM) and sulfolipids (2, 3). In recent years, many cell wall lipids have been found to be critical for various mycobacterial functions including biofilm formation (4–16). Biofilms are communities of organisms that are encased in a self-secreted extracellular matrix that can serve as a protective barrier against environmental and host derived threats (17). Perturbations in various classes of cell wall lipids have also been shown to yield altered biofilm phenotypes in various species of mycobacteria. Glycopeptidolipids (GPLs) are of notable interest as they are a major lipid class in the cell wall of non-tuberculous mycobacteria, and have been shown to play essential roles in the maintenance of cell wall integrity, colony morphology, surface motility, biofilm formation, antimicrobial susceptibility and host-pathogen interactions (11, 18–21). Strains defective in the synthesis and/or outer membrane transport of GPLs in various mycobacterial species were found to be defective in biofilm formation (10, 11, 22, 23). These findings suggest that lipid biosynthesis and transport pathways could serve as viable targets for the development of novel antibiofilm approaches against various mycobacterial species.

In this work, we further investigate the role of two conserved lipid transport pathways, LprG-Rv1410c and MmpL11, in cell wall biogenesis during biofilm formation in *Mycobacterium smegmatis* (*Msm*). We found that deletion of the *lprG*-*rv1410c* operon homologues *MSMEG_3070-3069* and *mmpL11* (*MSMEG_0241*) simultaneously led to similar biofilm defects as observed in the *MSMEG_3070-3069* and *mmpL11* mutants. This biofilm defect was found to be directly correlated with significant decreases in cellular c-di-GMP levels, but could also be due to changes in the synthesis and/or cell wall localization of glycopeptidolipids (GPLs), and changes in the expression of GPL biosynthesis and transport genes. Our findings therefore suggest that while both LprG-Rv1410c and MmpL11 pathways are involved in modulating cellular c-di-GMP levels during biofilm formation, only LprG-Rv1410c are important for regulating the synthesis and/or cell wall localization of GPLs, and MmpL11 may instead play a broad role in fine-tuning GPL levels during biofilm formation.

## RESULTS

### Loss of function in *MSMEG_3070-3069* or *mmpL11* overexpression correlates with decreased synthesis and/or transport of glycopeptidolipids

Although we previously found that the shared biofilm defect in the *MSMEG_3070-3069* or *mmpL11* mutant strains was not due to a shared change in the synthesis or localization of the biofilm and cell surface lipids: monomeromycolyl diacylglyceride (MMDAG), mycolate wax esters (MWE), triacylglycerides (TAG) and free mycolic acids (FMA) (24), we recently found that loss of function in both *MSMEG_3070-3069* and *mmpL11* simultaneously (Δ*MSMEG_3070-3069*/Δ*mmpL11*) had a biofilm defect that was similar to either the Δ*MSMEG_3070-3069* or Δ*mmpL11* mutant strains (Figure S1). This data supports a model in which LprG-Rv1410c and MmpL11 are likely acting in the same non-essential or parallel biochemical pathways to modulate biofilm formation by some other mechanism.

In recent years, glycopeptidolipids (GPLs), a cell surface specific lipid, has been found to be important for mycobacterial biofilm formation (10, 22, 25–28). Since we observed shared defects in biofilm formation for Δ*MSMEG_3070-3069*, Δ*mmpL11* and Δ*MSMEG_3070-3069*/Δ*mmpL11* mutant strains (Figure S1), we sought to determine if the shared biofilm defect between these strains could be connected to a change in the synthesis and/or localization of GPLs. To test this, we analyzed lipids extracted from the various mutant and wildtype *M. smegmatis* biofilms 5 days post inoculation via thin layer chromatography. Compared to wildtype, no change in GPL levels were observed in the total lipid extracts of the Δ*MSMEG_3070-3069::lprG-rv1410c*_*Mtb*_, Δ*mmpL11::mmpL11*_*Msm*_, or *mmpL11::Tn::mmpL11*_*Mtb*_ complement strains (Figure 1).

**Figure 1.**
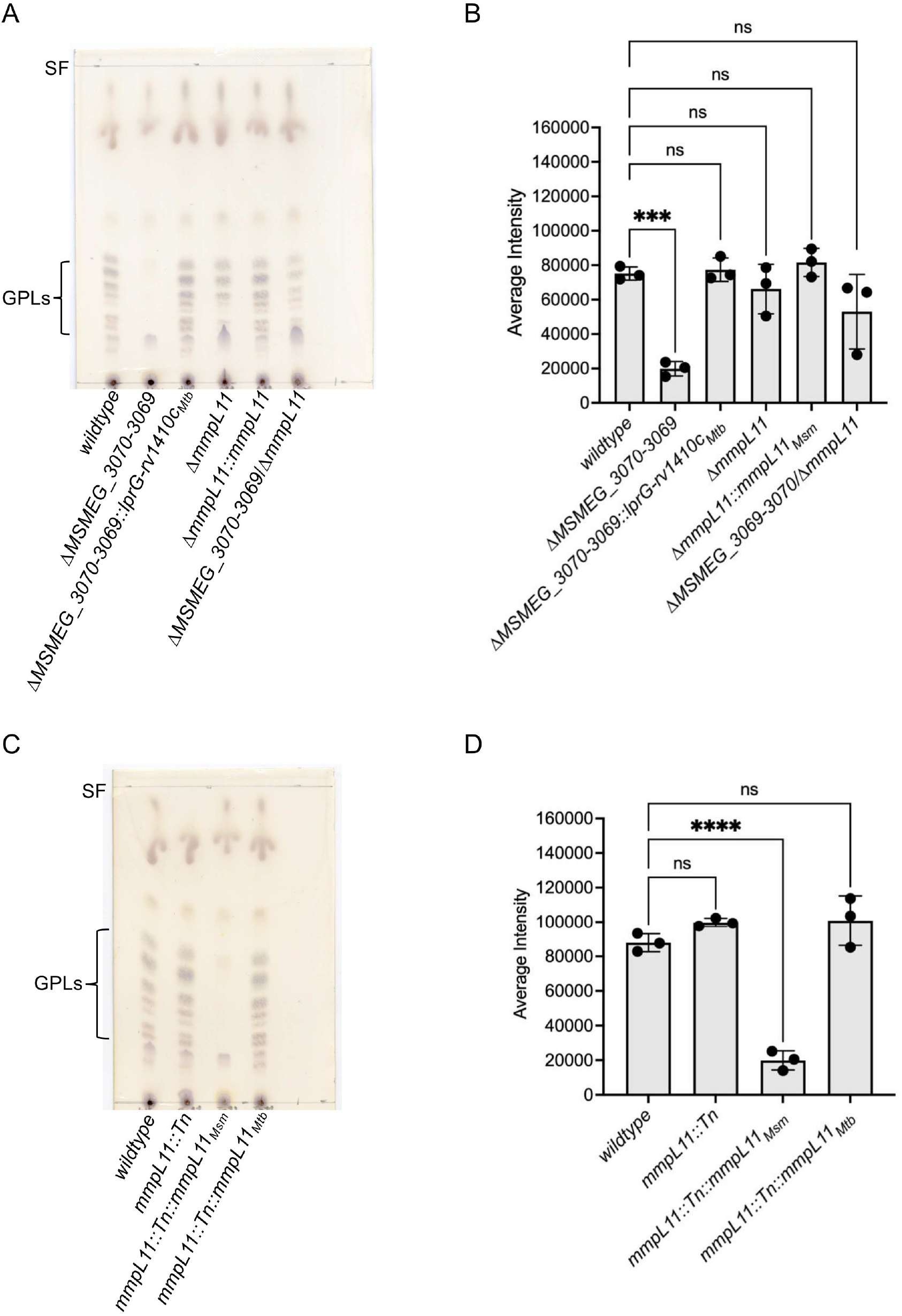
Glycopeptidolipid (GPL) profile was significantly altered in the total lipid extracts of the Δ*MSMEG_3070-3069*, and *mmpL11::Tn::mmpL11*_*Msm*_ strains. (**A**) TLC showing GPL levels in the parent wildtype, Δ*MSMEG_3070-3069*, Δ*MSMEG_3070-3069*/Δ*mmpL11*, Δ*mmpL11* and Δ*mmpL11::mmpL11* total lipid extracts. (**B**) Quantitative analysis of GPL levels in figure 1A. (**C**) TLC showing GPL levels in the parent wildtype, *mmpL11::Tn, mmpL11::Tn::mmpL11*_*Msm*_ *and mmpL11::Tn::mmpL11*_*Mtb*_ total lipid extracts. (**D**) Quantitative analysis of GPL levels in figure 1C. GPLs: glycopeptidolipids. SF: solvent front. 125 µg of total lipid extracts were resolved by TLC in chloroform:methanol (100:7, v/v) solvent system. GPLs were visualized by charring at 120°C after immersion in orcinol dip reagent (29). Data shown are the mean ± S.D. of three independent experiments. Statistical significance was determine using one-way ANOVA (Dunnett’s test) in GraphPad Prism version 11. (*\*\*\*\*, p= <0*.*0001* ; *\*\*\*, p= 0*.*0002* ; *\*\*, p= 0*.*0021*; *\*, p= 0*.*03*; ns: not significant).

Consistent with previous research work (7), no change in GPL levels were observed in the total lipid extracts of the *mmpL11::Tn* mutant strain (Figures 1A and1B). Excitingly, we observed a complete abrogation of GPLs in the total lipid extracts of the Δ*MSMEG_3070-3069* mutant and *mmpL11::Tn::mmpL11*_*Msm*_ complement strain that were statistically significant (Figures 1A, 1B, 1C and 1D). We previously found that mmpL11 is over expressed in the *mmpL11::Tn::mmpL11*_*Msm*_ complement strain (∼64 fold increase compared to the wildtype) (24). Further, we observed a slight decrease (although not statistically significant) in GPL levels in the total lipid extracts of the ΔmmpL11 and Δ*MSMEG_3070-3069*/Δ*mmpL11* strains (Figures 1A and 1B). These findings are in contrast to our mmpL11::Tn mutant strain data (Figures 1C and 1D) and previously published work (7), but nevertheless suggest that LprG-Rv1410c and MmpL11 are likely involved in regulating GPL biosynthesis and/or transport.

### Loss of *MSMEG_3070-3069* or *mmpL11 overexpression* correlates with decreased expression of GPL synthesis and surface localization genes

We next sought to determine if the observed changes in GPL levels in the total lipid extracts could be connected to changes in gene expression of GPL biosynthesis and/or surface localization genes. To test this, RNA sequencing was performed on total RNA extracted from select mutant and wildtype *M. smegmatis* biofilms 5 days post inoculation. Consistent with our thin-layer chromatography (TLC) data, we observed no changes in gene expression of the GPL biosynthesis or surface localization genes in the Δ*mmpL11*, Δ*MSMEG_3070-3069*/Δ*mmpL11* and *mmpL11::Tn* mutant strain (Figures 2B, 2C and 2D). Compared to wildtype and consistent with our TLC data, however, we found that the GPL biosynthesis and surface localization genes *MSMEG_0400, MSMEG_0402, MSMEG_0403, MSMEG_0404* and *MSMEG_0405* (21, 30, 31) were significantly downregulated in the Δ*MSMEG_3070-3069* mutant and *mmpL11::Tn::mmpL11*_*Msm*_ complement strains (Figures 2A and 2E). Our findings further underscore a role for LprG-Rv1410c and MmpL11 in regulating GPL biosynthesis and/or transport during *M. smegmatis* biofilm formation.

**Figure 2.**
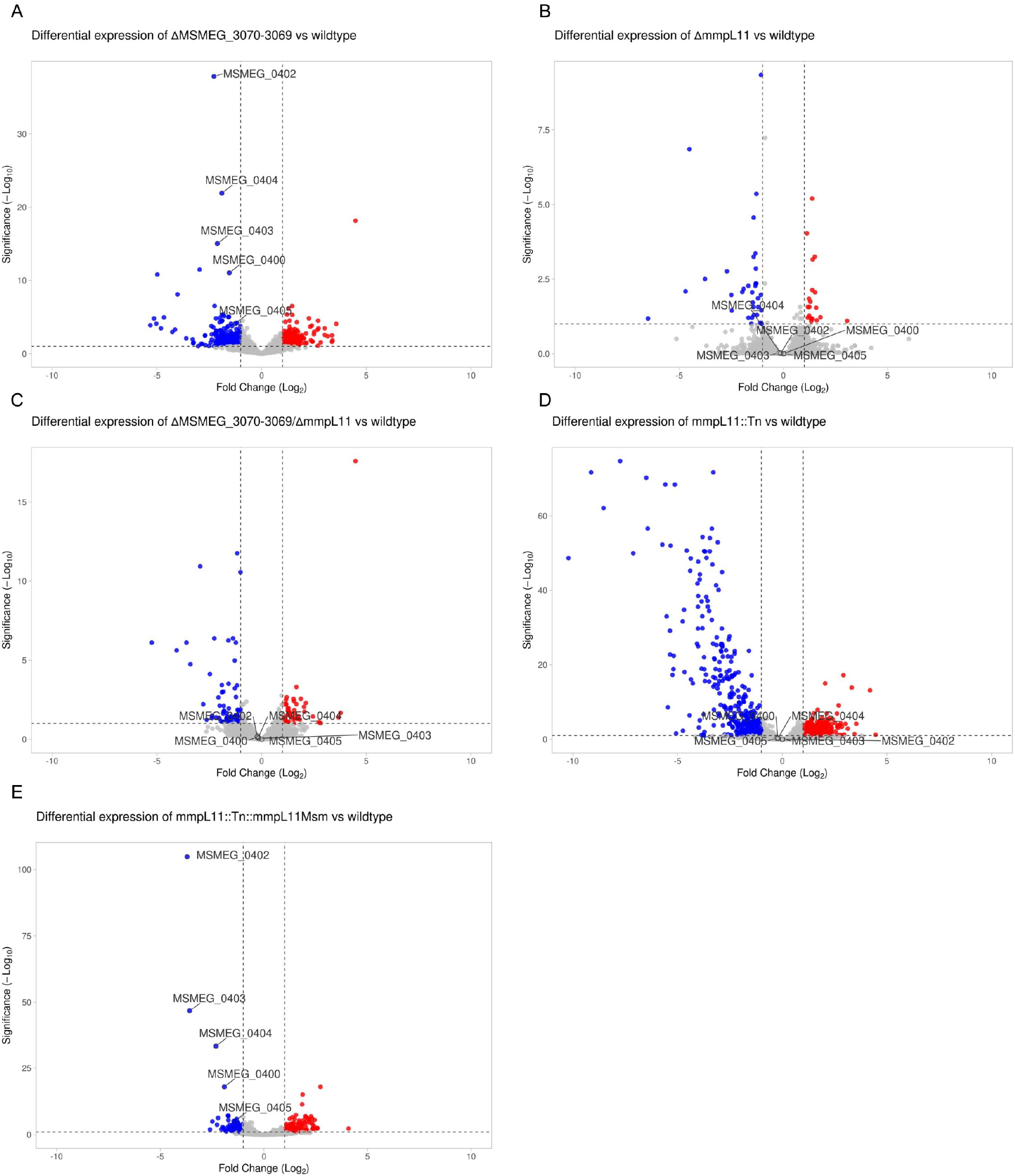
Glycopeptidolipid (GPL) biosynthesis and/or surface localization genes were significantly downregulated in the Δ*MSMEG_3070-3069* and *mmpL11::Tn::mmpL11*_*Msm*_ strains. (**A**) Differential expression of Δ*MSMEG_3070-3069* compared to wildtype. (**B**) Differential expression of Δ*mmpL11* compared to wildtype. (**C**) Differential expression of Δ*MSMEG_3070-3069*/Δ*mmpL11* compared to wildtype. (**D**) Differential expression of *mmpL11::Tn* compared to wildtype. (**E**) Differential expression of *mmpL11::Tn::mmpL11*_*Msm*_ compared to wildtype. Genes with *p<0*.*05* and a log_2_ (fold-change)≥1 (upregulated) or ≤-1(downregulated) are shown. Blue dots represent downregulated genes, red dots represent upregulated genes and grey dots represent genes with no changes in expression. Data shown are representative of five independent experiments. Volcano plots were generated using VolcaNoseR (32).

### Loss of *MSMEG_3070-3069* and/or *mmpL11* as well as overexpression of *mmpL11* correlates with decreased cellular levels of c-di GMP during biofilm formation

Bis-(3′-5′)-cyclic dimeric GMP (c-di-GMP) is a secondary messenger molecule that has been demonstrated to modulate biofilm formation in many groups of bacteria (33, 34) including mycobacteria (12, 13, 35–37). Further, c-di-GMP is thought to be involved in regulating GPL metabolism in *Mycobacterium smegmatis* (35). Given that we observed a shared biofilm defect in the Δ*MSMEG_3070-3069*, Δ*mmpL11*, Δ*MSMEG_3070-3069*/Δ*mmpL11* and *mmpL11::Tn::mmpL11*_*Msm*_ strains, and found altered GPL levels in the Δ*MSMEG_3070-3069* and *mmpL11::Tn::mmpL11*_*Msm*_ strains, we reasoned that one or both of these phenotypes may be correlated with altered cellular c-di-GMP levels. Indeed, compared to wildtype, we observed a significant decrease in cellular c-di-GMP levels in the Δ*MSMEG_3070-3069*, Δ*mmpL11*, Δ*MSMEG_3070-3069*/Δ*mmpL11, mmpL11::Tn* and *mmpL11::Tn::mmpL11*_*Msm*_ strains (Figures 3A and 3B).

**Figure 3.**
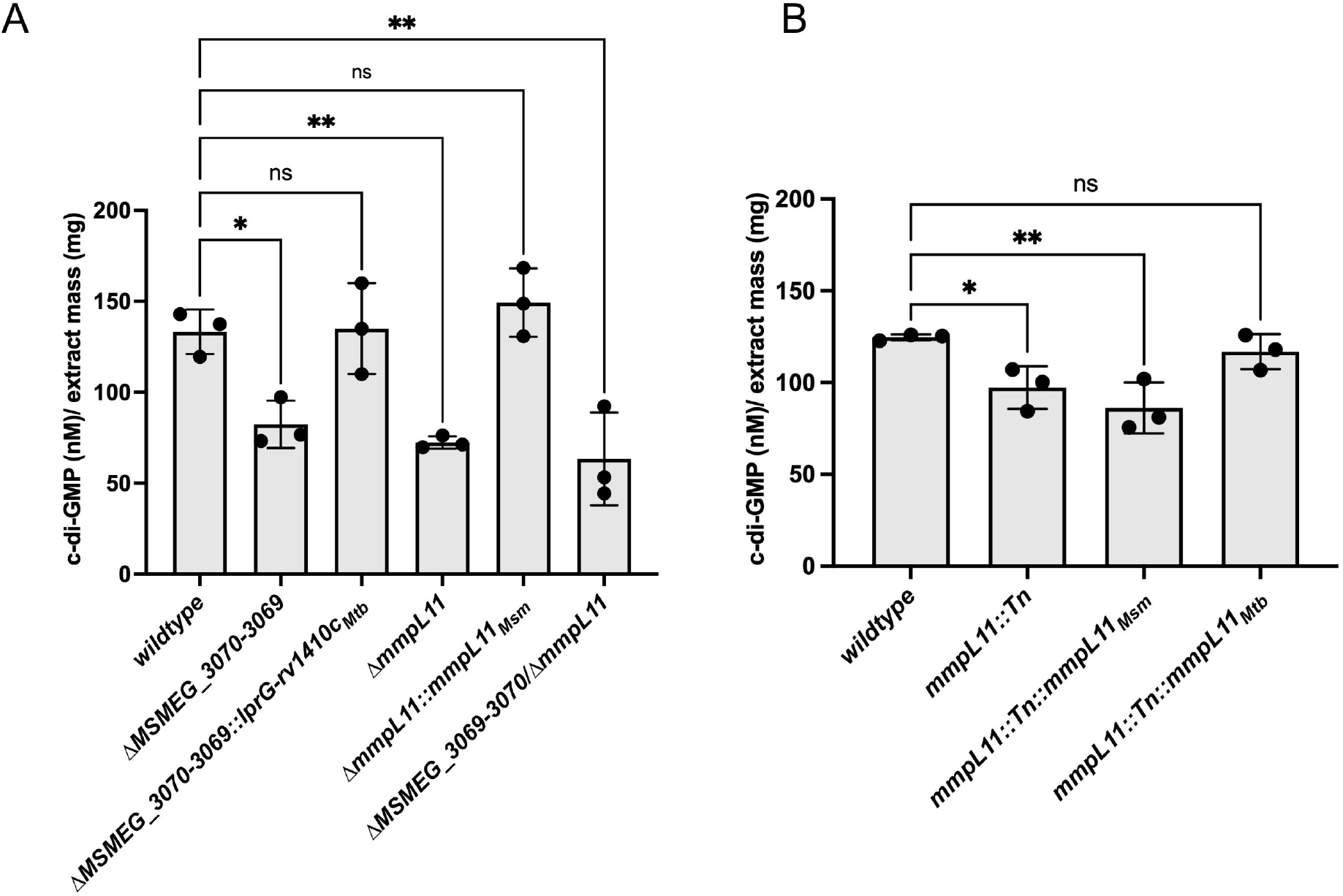
Cellular c-di-GMP levels were significantly decreased in the Δ*MSMEG_3070-3069*, Δ*mmpL11*, Δ*MSMEG_3070-3069*/Δ*mmpL11, mmpL11::Tn* and *mmpL11::Tn::mmpL11*_*Msm*_ strains. (**A**) Quantitative analysis of cellular c-di-GMP levels of parent wildtype, Δ*MSMEG_3070-3069*, Δ*MSMEG_3070-3069*/Δ*mmpL11*, Δ*mmpL11* mutant and associated complement strains. (**B**) Quantitative analysis of cellular c-di-GMP levels of parent wildtype, Δ*MSMEG_3070-3069*, Δ*MSMEG_3070-3069*/Δ*mmpL11*, Δ*mmpL11* mutant and associated complement strains. parent *mmpL11::Tn* mutant and associated complement strains. Data shown are the mean ± S.D. of three independent experiments. Statistical significance was determine using one-way ANOVA (Dunnett’s test) in GraphPad Prism version 11. (*\*\*, p= 0*.*0021*; *\*, p= 0*.*03*; ns: not significant).

Wildtype-like c-di-GMP levels, however, were observed for the *mmpL11::Tn::mmpL11*_*Mtb*_, Δ*MSMEG_3070-3069* and Δ*mmpL11* complement strains (Figures 3A and 3B). These findings suggest that LprG-Rv1410c and MmpL11 are important for modulating cellular levels of c-di-GMP during biofilm formation.

## DISCUSSION

In this work, we demonstrate that LprG-Rv1410c (*MSMEG_3070-3069*) and MmpL11 likely act in non-essential parallel biochemical pathways to regulate biofilm formation in *M. smegmatis* via modulation of GPL and cellular c-di-GMP levels. We found that loss of function in *MSMEG_3070-3069* and mmpL11 simultaneously (all three genes) led to GPL levels that were more similar to *mmpL11* levels compared to wildtype or *MSMEG_3070-3069*, suggesting that *MSMEG_3070-3069* may be important for regulating the biosynthesis and/or surface localization of GPLs, but *mmpL11* may instead be important for fine tuning GPL levels overall. Both the *MSMEG_3070-3069* mutant and the *mmpL11::Tn::mmpL11*_*Msm*_ complement strains had complete abrogations of GPL levels in the total lipid extracts compared to the wildtype, Δ*mmpL11*, Δ*MSMEG_3070-3069*/Δ*mmpL11* or *mmpL11::Tn* strains suggesting that loss of function in *MSMEG_3070-3069* or overexpression of *mmpL11* led to perturbations in the biosynthesis or surface localization of GPLs, which in turn led to significant decreases in GPL levels and subsequent disruptions in pellicle biofilm formation. Although we only observed slight decreases in GPL levels in the *MSMEG_3070-3069/mmpL11* and *mmpL11* mutants, it is probable that these slight alterations in GPLs could have still led to disruptions in biofilm formation as perhaps only small changes in GPLs are necessary for alterations in biofilm formation to occur. Furthermore, given that TLC can only resolve major lipid classes from each other but cannot provide detailed information on subclasses of a specific lipid type, it is also possible that certain subclasses of GPLs could be altered in both the *MSMEG_3070-3069/mmpL11* and *mmpL11* mutants, thus yielding biofilm defects that were comparable to the *MSMEG_3070-3069* mutant and the *mmpL11::Tn::mmpL11*_*Msm*_ complement strains, and a slight decrease in the GPL profile overall. Further experiments, however, are needed to determine if one or both of these phenomena underlie the observed shared biofilm defect in the *MSMEG_3070-3069/mmpL11* and *mmpL11* mutants.

We hypothesized that the perturbations in GPL levels could be due to changes in expression of GPL biosynthesis and/or surface location genes. Consistent with our TLC data, we did observe that GPL biosynthesis and surface localization genes were downregulated in the *MSMEG_3070-3069* mutant and the *mmpL11::Tn::mmpL11*_*Msm*_ complement strains, suggesting that the observed decrease in GPL levels in the total lipid extracts were indeed due to decreased expression of GPL biosynthesis and surface localization genes. Further, the slight decrease in GPLs (and therefore the defect in biofilm formation) in the *MSMEG_3070-3069*/*mmpL11* and *mmpL11* mutants, may instead be due to decrease in activity of the GPL biosynthesis and/or surface localization proteins rather a change in gene expression. Additional experiments, however, are needed to determine if this is indeed the case.

Finally, given that c-di-GMP is an established regulator of biofilm formation (33, 34) and is thought to regulate GPL metabolism in *M. smegmatis* (35), we sought to determine if the shared biofilm defect between the various strains could be connected to altered cellular levels of c-di-GMP during biofilm formation. We found that there was indeed a significant decrease in cellular c-di-GMP levels in the *MSMEG_3070-3069, mmpL11, MSMEG_3070-3069*/*mmpL11, mmpL11::Tn* mutant strains and *mmpL11::Tn::mmpL11*_*Msm*_ complement strain compared to wildtype, suggesting that both the *MSMEG_3070-3069* and *mmpL11* pathways are important for modulating c-di-GMP levels during biofilm formation. Although we did not observe a change in gene expression for *dcpA* (Tables S3-S7), the gene that encodes for the bifunctional protein DcpA which synthesizes and hydrolyses c-di-GMP in *M. smegmatis* (38–40), it is still probable that a change in DcpA activity could have occurred, which in turn led to decreased levels of c-di-GMP and subsequent biofilm defects in the aforementioned strains. How changes in cellular c-di-GMP levels specifically modulate changes in GPL levels is still unclear at this time, however, as c-di-GMP effectors that regulate the expression of GPL biosynthesis and/or surface localization genes are yet to be identified.

In summary, our research findings collectively suggest LprG-Rv1410c may be directly involved in GPL biosynthesis and/or surface localization such that loss of function in these genes led to a complete abrogation in GPL levels in the total lipid extracts of the Δ*MSMEG_3070-3069* strain. MmpL11, however, may play a role in broadly fine-tuning GPL levels as overexpression of the MmpL11 in the *mmpL11::Tn::mmpL11*_*Msm*_ strain also led to a complete abrogation of GPL levels. Conversely, loss of function or disruption of MmpL11 in the Δ*MSMEG_3070-3069*/Δ*mmpL11*, Δ*mmpL11* or *mmpL11::Tn* strains respectively, led to slight but not significant changes in GPL levels overall.

## MATERIALS AND METHODS

### Bacterial strains, Culture media and Culture conditions

Bacterial strains used in this study are listed in Table S1. *Mycobacterium smegmatis* mc^2^155 served as the parent (wild-type) strain. For planktonic growth, *M. smegmatis* was cultured at 37°C with agitation at 250 rpm in Middlebrook 7H9 broth supplemented with 10% albumin/dextrose/catalase (ADC), 0.5% glycerol, and 0.05% Tween 80 (all % are *v/v* unless otherwise indicated). For growth on solid medium, unless otherwise indicated, *M. smegmatis* was plated on Middlebrook 7H11 agar containing 10% ADC, 0.5% glycerol, and 0.05% Tween 80. When required, hygromycin, kanamycin and/or zeocin were added to the growth medium at 50, 25, or 10 µg/mL, respectively.

### Construction of Mutant and Complement *M. smegmatis* strains

All primers and plasmids used in the construction of mutant and complement strains are listed in Table S2. The Δ*MSMEG_3070-3069*, Δ*MSMEG_3070-3069::lprG-rv1410c*_*Mtb*_ and wildtype parent *M. smegmatis* strains were a gift from Eric Rubin (41, 42). The *mmpL11::Tn, mmpL11::Tn::mmpL11*_*Msm*_, *mmpL11::Tn::mmpL11*_*Mtb*_ and wildtype parent *M. smegmatis* strains were a gift from Georgiana Purdy (7). A Δ*MSMEG_3070-3069*/Δ*mmpL11* strain was generating via recombineering (43) using the Δ*MSMEG_3070-3069* strain as the genetic background using previously described methods (24).

### Biofilm Growth

For biofilm assays, *M. smegmatis* strains were cultured as previously described (7). Briefly, *M. smegmatis* was inoculated at OD_600_ 0.05 in Sauton’s medium, without Tween 80, in polystyrene Petri dishes (100 mm × 15 mm) and incubated at 30°C without disturbance for up to 5 days. Sauton’s medium contained 0.5 g/L monobasic potassium phosphate, 0.5 g/L anhydrous magnesium sulfate, 4.0 g/L L-asparagine, 0.05 g/L ferric ammonium citrate, 2.0 g/L anhydrous citric acid, 4.76% glycerol, and 1 mg/L zinc sulfate heptahydrate at pH 7.0.

### Lipid extraction and Analysis

For total lipid extraction, pellicle biofilms were harvested 5 days after inoculation via centrifugation at 3000× *g* for 10 min, the supernatants were discarded and the biofilm pellets were subsequently resuspended in 5 mL chloroform/methanol (2:1, *v*/*v*) and vortexed for 30 s. The extracts were then clarified via centrifugation at 1000× *g* for 10 min and the supernatants were dried under nitrogen gas at 30°C (Biotage TurboVap LV).

For analysis via thin-layer chromatography, total lipid extracts were resuspended in chloroform/methanol (2:1, *v*/*v*) and spotted onto silica plates (Millipore-Sigma, Chicago, IL, USA 1.05553.0001). Loads were normalized according to dry weight. Glycopeptidolipids (GPLs) were resolved with chloroform/methanol (100:7, *v/v*). The total amount of GPLs were calculated via densitometry using ImageJ (44), and the mean ± standard deviation (S.D.) from three independent experiments were analyzed. Statistical analysis was performed using GraphPad Prism version 11.

### C-di-GMP Extraction and Quantification

5 days post inoculation, formaldehyde was added to a final concentration of 0.18% to the pellicle biofilms which were then harvested via centrifugation at 3000 *xg* for 10 min at 4 °C. The supernatants were then discarded, the pellets were resuspended in 300 µL ice cold lysis buffer (40% acetonitrile, 40% methanol, 0.1% formic acid and 19.9% millique grade water), heated at 95 °C for 3 mins and then cooled on ice for 5 min. The samples were then centrifuged at 3000 *xg* for 10 min at 4 °C to remove insoluble material, and the supernatant was subsequently transferred to new microcentrifuge tubes. Thereafter, the resulting pellets were extracted twice with 300 µL of extraction buffer at 4 °C omitting the heating step. The supernatants of the three extractions were combined and dried using a centrifugal evaporator. The c-di-GMP content of the dried extracts was quantified using the Lucerna c-di-GMP assay kit (Lucerna Technologies, Brooklyn, NY, USA) as per the manufacturer’s instructions. Total c-di-GMP content was normalized according to dry extract mass.

### RNA Extraction, RNA sequencing and RNA-seq data processing and analysis

Pellicle biofilms were harvested 5 days after inoculation via centrifugation at 3000 *xg* for 10 min and the pellets were resuspended in 1 mL TRIzol LS (Thermo Fisher Scientific, Waltham, MA, USA) and stored at -80 °C. TRIzol resuspensions were thawed on ice and subsequently lysed via bead beating (BeadRupter 12, OMNI International, Kennesaw, GA, USA) using 0.1 mm zirconia/silica beads (Biospec, Bartlesville, OK, USA) for 30 s at 6 m/s followed by 5 min incubation on ice for a total of 4 cycles. The lysates were clarified via centrifugation at 12,000 *xg* at 4 °C for 10 min and total RNA was extracted and purified from the aqueous phase using chloroform/isoamyl alcohol (24:1, v/v) and the Qiagen Rneasy kit, respectively. RNAseOut (Thermo Fisher Scientific, Waltham, MA, USA) was added to prevent RNA degradation. DNA contamination was removed using the TurboDNAse free kit, (Thermo Fisher Scientific, Waltham, MA, USA) and the concentration and quality of RNA (260 nm/280 nm absorbance) was assessed using a nanodrop spectrophotometer. Samples were then stored at -80 °C until submission to SeqCenter (Pittsburg, PA, USA).

Demultiplexing, quality control library preparation, ribodepletion, adapter trimming and paired-end sequencing were performed on the NovaSeq X Plus by SeqCenter. Raw read quality was assessed using FastQC (version 0.12.1). Sequencing reads were mapped to the *Mycobacterium smegmatis mc*^*2*^*155* strain genome (RefSeq assembly accession: NC_008596.1) using HISAT2 (version 2.2.1) with the default paired-end parameters and the very sensitive flag. Gene counts were filtered and quantified using samtools (version 1.20) and featureCounts (version 2.1.1). Read count normalization and differential expression analysis were performed using DESeq2 (version 1.38.3). Genes were considered to be differentially expressed when the adjusted p-value > 0.05 and the log_2_-fold change) ≥1 (upregulated) or ≤-1(downregulated). Volcano plots were generated using VolcaNoseR (32).

## ACKNOWLEDGEMENTS

We thank Eric Rubin for providing the wildtype, Δ*MSMEG_3070-3069* and Δ*MSMEG_3070-3069::lprG-rv1410c*_*Mtb*_ strains. We thank Georgiana Purdy for providing the wildtype, *mmpL11::Tn, mmpL11::Tn::mmpL11*_*Msm*_ and *mmpL11::Tn::mmpL11*_*Mtb*_ strains. We thank Jeffrey Cox for providing the pJSC407 plasmid. We thank Christopher Sassetti for providing the pNit-RecET-SacB-Kan plasmid. We thank Anil Ojha for his feedback and helpful discussions.

## DATA AVAILABILITY

RNAseq data is available on NCBI GEO accession number GSE346443

## SUPPLEMENTARY MATERIAL

Figure S1: 5 day time course of pellicle biofilm formation at the air-liquid interface.

Table S1: Strains used in this study

Table S2: Plasmids and primers used in this study

Table S3: Differential expression analysis of *MSMEG_3070-3069* mutant and wildtype

Table S4: Differential expression analysis of *mmpL11* mutant and wildtype

Table S5: Differential expression analysis of Δ*MSMEG_3070-3069*/Δ*mmpL11* mutant and wildtype

Table S6: Differential expression analysis of *mmpL11::Tn* strain and wildtype

Table S7: Differential expression analysis of *mmpL11::Tn::mmpL11*_*Msm*_ strain and wildtype

