## Supplementary information for "Loss of function of LprG-Rv1410c and MmpL11 homologues in *Mycobacterium smegmatis* leads to altered glycopeptidolipid profile and decreased cellular c-di-GMP levels during biofilm formation"

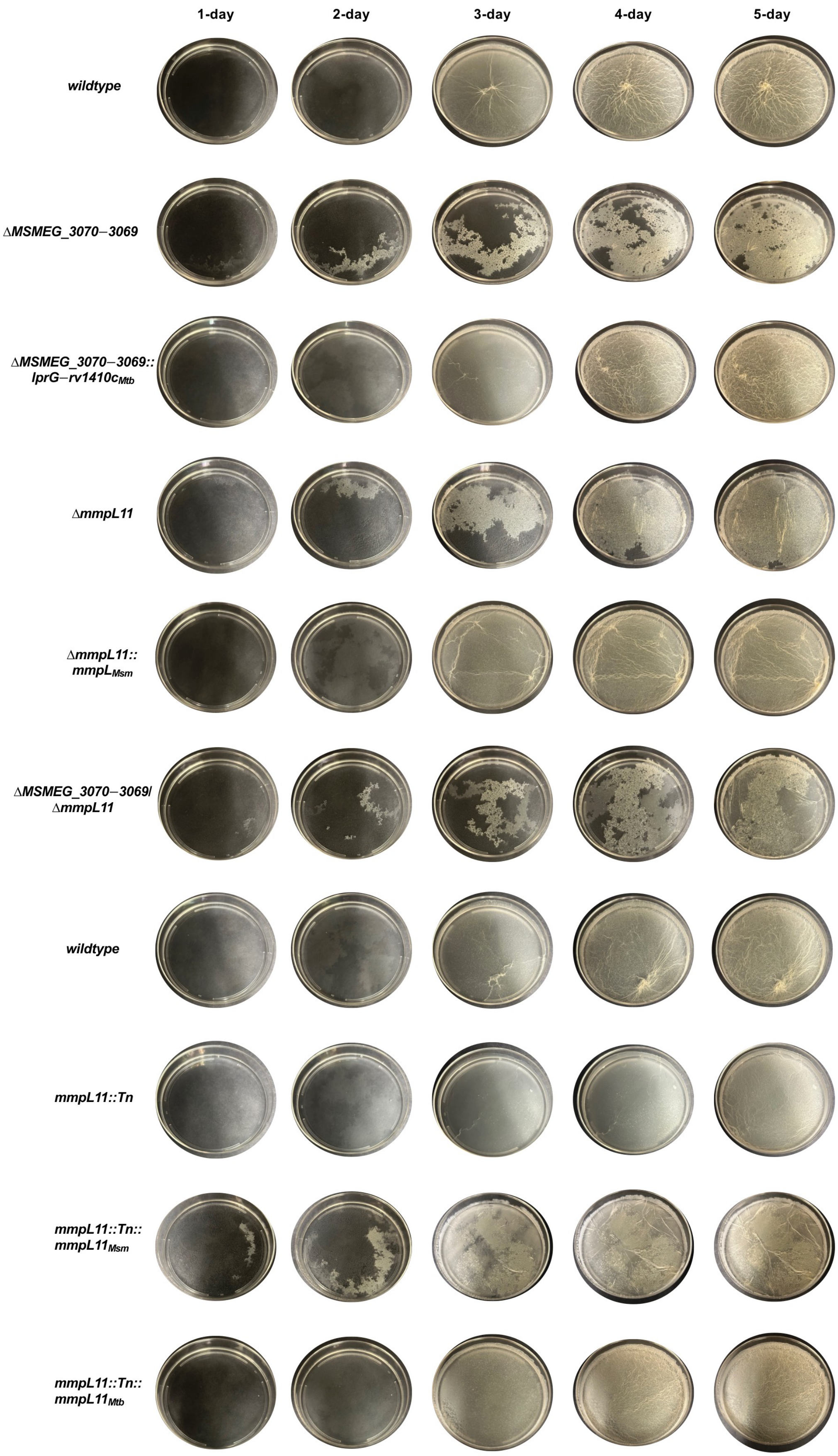
1-day
2-day
3-day
4-day
5-day
wildtype
ΔMSMEG_3070−3069
ΔMSMEG_3070−3069::
lprG−rv1410c$_{Mtb}$
ΔmmpL11
ΔmmpL11::
mmpL$_{Msm}$
ΔMSMEG_3070−3069/
ΔmmpL11
wildtype
mmpL11::Tn
mmpL11::Tn::
mmpL11$_{Msm}$
mmpL11::Tn::
mmpL11$_{Mtb}$

Figure S1 5 day time course of pellicle biofilm formation at the air-liquid interface. Equal number of bacteria were inoculated in Sauton's medium without Tween 80 in polystyrene petri dishes and the plates were incubated at 30°C without disturbance for 5 days. Δ*MSMEG_3070−3069*, Δ*mmpL11*, Δ*MSMEG_3070−3069*/Δ*mmpL11* and *mmpL11::Tn::mmpL11*$_{Msm}$ strains were defective in pellicle biofilm formation starting at the day 2 time point. *mmpL11::Tn* was defective in pellicle biofilm formation at the day 4 time point. The data shown is representative of five biological replicates.

Table S1 Strains used in this study

| Species | Parent strain | Integration | Insert | Marker | Source |
|---|---|---|---|---|---|
| Mycobacterium smegmatis mc$^2$155 | | | | | ATCC 700084 |
| | Δ*MSMEG_3070−3069* | none | none | none | (1) |
| | Δ*MSMEG_3070−3069* | L5 attB | Phsp70−*lprG*−*rv1410c* | Zeo | (2) |
| | Δ*MSMEG_0241* | none | none | Hyg | (3) |
| | Δ*MSMEG_0241* | L5 attB | Pnative−*MSMEG_0241* | Hyg/Kan | (3) |
| | Δ*MSMEG_3070−3069*/Δ*MSMEG_0241* | none | none | Hyg | This work |
| | *MSMEG_0241::Tn* | none | pVV16 | Kan | (4) |
| | *MSMEG_0241::Tn* | none | pVV16−*MSMEG_0241* | Kan | (4) |
| | *MSMEG_0241::Tn* | L5 attB | Pnative−*mmpL11* | Hyg/Kan | (4) |

Zeo: Zeocin; Hyg: Hygromycin B; Kan: Kanamycin

Table S2 Plasmids and Primers used in this study

| Plasmid | Description |
|---|---|
| pJSC407 | Mycobacteria knockout plasmid; Hyg resistance |
| pmlp082 | pJSC407 with 125bp upstream and downstream fragments of *mmpL11* gene; Hyg resistance |
| pNIT-RecEt-SacB-Kan | Plasmid with Che9c RecET gene for recombineering in mycobacteria; Kan resistance |
| pMV306 | Single copy integrating plasmid with L5 integrase; integrates at mycobacteria chromosomal *attB* site; Kan resistance |
| pmlp083 | pMV306 with mmpL11 inserted into XbaI/ClaI sites; Kan resistance |
| **Primer** | **Sequence** |
| **Gene deletion** | |
| omlp741 | TGGATCCACGAAGCTTTGGTCAGAGCCTGGTTGGTC |
| omlp742 | GGCCACCATGAAGCTTCTACAAGCGCATCATGAACTGCTGGATG |
| omlp743 | CGGACAGGACTCTAGACTGGAGGAGGCGAAGTGACG |
| omlp744 | CCGGGGATCCTCTAGAGCACGAGAACTTCCGACAG |
| **Complementation** | |
| omlp745 | GATCTTTAAATCTAGAGTGTCCAGTTTCTTGCCTTGC |
| omlp746 | ACTACGTCGACATCGATTCACTTCGCCTCCTCCAGCATTG |
| **PCR screening** | |
| ojcs240 | CAGGCTCGCGTAGGAATCATC |
| **Sequencing** | |
| oevv137 | GATGGCATAAAACGAAAGGCC |
| ojcs238 | GCCTTTGAGTGAGCTGATACC |
| omlp336 | CGCCTGATCAGGATCGTAATAC |
| omlp337 | GACCTCGACGACCTGCAG |
| omlp338 | CTGACGCTCAGTGGAACG |
| omlp051 | CATCTTCGTGGACCTGGCC |
| omlp756 | CTACCTCGCTCTCAACCAGTC |
| omlp757 | GTCACCGGCATCTACCTCATC |
| omlp758 | GATCAGCTCCCTGGACA |
